# Clinical metagenomics of bone and joint infections: a proof of concept study

**DOI:** 10.1101/090530

**Authors:** Etienne Ruppé, Vladimir Lazarevic, Myriam Girard, William Mouton, Tristan Ferry, Frédéric Laurent, Jacques Schrenzel

## Abstract

**Background:** Bone and joint infections (BJI) are severe infections that require a tailored and protracted antibiotic treatment. The diagnostic of BJI relies on the culture of surgical specimens, yet some bacteria would not grow because of extreme oxygen sensitivity or fastidious growth. Hence, metagenomic sequencing could potentially address those limitations. In this study, we assessed the performances of metagenomic sequencing of BJI samples for the identification of pathogens and the prediction of antibiotic susceptibility.

**Methods:** A total of 179 samples were considered. The DNA was extracted with a kit aiming to decrease the amount of human DNA (Molzym), and sequenced on an Illumina HiSeq2500 in 2x250 paired-end reads. The taxonomy was obtained by MetaPhlAn2, the bacterial reads assembled with MetaSPAdes and the antibiotic resistance determinants (ARDs) identified using a database made of Resfinder+ARDs from functional metagenomic studies.

**Results:** We could sequence the DNA from 24 out of 179 samples. For monomicrobial samples (n=8), the presence of the pathogen was confirmed by metagenomics in all cases. For polymicrobial samples (n=16), 32/55 bacteria (58.2%) were found at the species level (41/55 [74.5%] at the genus level). Conversely, a total of 273 bacteria not found in culture were identified, 182 being possible pathogens undetected in culture and 91 contaminants. A correct antibiotic susceptibility could be inferred in 94.1% cases for monomicrobial samples and in 76.5% cases in polymicrobial samples.

**Conclusions:** When sufficient amounts of DNA can be extracted from samples, we found that clinical metagenomics is a potential tool to support conventional culture.

## Introduction

Bone and joint infections (BJI) are severe infections that affect a growing number of patients [1]. Along with the surgical intervention, the microbiological diagnosis is a keystone of the management of BJI in *(i)* identifying the bacteria causing the infection and (*ii*) assessing their susceptibility to antibiotics. Currently, this is achieved by culturing surgical samples on various media and conditions, together with a long time of incubation to recover fastidiously-growing bacteria that can be involved in BJI. Still, some bacteria would not grow under these conditions because of extreme oxygen sensitivity, a prior antibiotic intake or metabolic issues (quiescent bacteria in chronic infections). Consequently, the antibiotic treatment may not span all the bacteria involved in the infection, which can favor the relapse and the need for a new surgery.

Clinical metagenomics refers to the concept of sequencing all the DNA (i.e. all the genomes) present in a clinical sample with the purpose of recovering pathogens and inferring their antibiotic susceptibility pattern [2]. This new, culture-independent method takes advantages of the thrilling development of next-generation sequencing (NGS) technologies since the mid-2000s. These sequencers typically yield thousands to millions of short reads (sequences of size ranging from 100 bp to a few kbp), which virtually enables to recover the sequences of all the genes present in the sample, yet in a disorganized fashion. Substantial bio-informatics efforts are thereby needed to re-construct and re-order the original sequences in genomes, and are referred to as the assembly process. Hence, various information such as the taxonomic identification of the present species, antibiotic resistance determinants (ARDs), mutations (as compared to a reference genome or sequence), single nucleotide polymorphisms (for clonality assessment) and virulence genes can be found.

Clinical metagenomics is an emerging field in medicine. So far, a few attempts to use metagenomics on clinical samples have been performed (on urines [3,4], cerebrospinal fluid or brain biopsy [5,6], blood [7] and skin granuloma [8]) likely because of the high price of metagenomics and the complexity of the management of sequence data for clinical microbiologists. To the best of our knowledge, metagenomics has never been applied to BJI samples.

As for the inference of antibiotic susceptibility testing from the genomic information, a few studies focusing on *Escherichia coli*, *Klebsiella pneumoniae*, *Mycobacterium tuberculosis* and *Staphylococcus aureus* have constantly showed excellent correlations between the analysis of the genomic content of antibiotic resistance determinants (ARDs) and the phenotype [9–15] while performances were not as good for *Pseudomonas aeruginosa* [16]. Furthermore in metagenomic data, the possible presence of multiple pathogens poses the issue of linking ARDs to their original host in order to infer its antibiotic susceptibility pattern [3]. So far no method has been proposed to address this question. Applying metagenomics in the context of BJI is thus seducing in that 1) there is no limit in the number of species and ARDs that could be detected (as opposed to PCR-based methods which detect targeted bacteria), 2) unculturable bacteria, fastidious growers (such as *Propionibacterium* sp.) or bacteria altered by prior antibiotic use would be recovered, and 3) the antibiotic susceptibility inference would benefit from both the detection of ARDs (such as *mecA*, *qnr, dfr, erm, etc.*) and the identification of mutations leading to resistance to key antibiotics used in BJI. Here, our main objective is to assess the performances of clinical metagenomics in BJI in terms of pathogen identification and inference of AST, as compared with conventional microbiology (gold standard).

## Material and methods

### Samples

We initially included 179 per-operative samples recovered from 47 patients (range 1-8 samples per patient). All but 2 (swabs) were solid specimens. The quantity of material for each non-swab sample (n=177) was macroscopically estimated: less than 1 mL (n=100), from 1 to 10 mL (n=60) and more than 10 mL (n=17). The samples were collected from September 2015 to January 2016 in the orthopedic departments of the CRIOAc (Regional Reference Center for Complex Osteo-Orticular Infections), Lyon, France (https://www.crioac-lyon.fr) and stored at –80ºC until shipment in dry ice to the Genomic Research Laboratory in Geneva on April 13, 2016. The samples had previously been cultured (see Supplementary Methods for the detailed protocol): a single bacteria or yeast was recovered for 104 out of 179 samples (58.1%), the remaining yielding 2 (24/179, 13.4%), 3 (26/179, 14.5%), 4 (14/179, 7.8%) or 5 (11/179, 6.1%) bacteria and yeasts. The exploitation of the collection used in this study was approved by the Ethical Committee of the Lyons University Hospital (September 25, 2014).

### DNA manipulations

The detailed protocol for DNA manipulations can be found in the Supplementary Methods. Briefly, the DNA from samples was extracted by the Ultra-Deep Microbiome Prep kit (Molzym, Bremen, Germany) according to the manufacturer’s instructions (Version 2.0) for tissue samples. The concentration of bacterial and human DNA was determined by qPCR experiments as described previously [17]. About 3 ng of DNA were sent to Fasteris (Plan-les-Ouates, Switzerland) for DNA purification and subsequent sequencing in Rapid Run mode for 2×250+8 cycles on an Illumina HiSeq 2500 instrument (with a HiSeq Rapid Flow Cell v2).

### Bioinformatic methods

The pipeline of read processing is displayed in Figure 1, and detailed in the Supplementary methods. Briefly, quality-filtered reads were processed with MetaPhlAn2 to get the taxonomic profile of the microbial community [18,19]. The bacterial reads assembled using metaSPAdes [20]. The identification of ARDs was achieved in mapping the total quality-filtered reads onto a database made of the ARDs from the ResFinder database [21] and ARDs from functional metagenomic studies [22–24]. To get the depth of sequencing of the bacterial species and of the ARDs in samples, and the single nucleotide variants (SNVs), we separately mapped the reads against the contigs assigned to one given species and against the ARDs identified in this sample The same pipeline was applied to all samples after downsizing to 1M reads.

**Figure 1:**
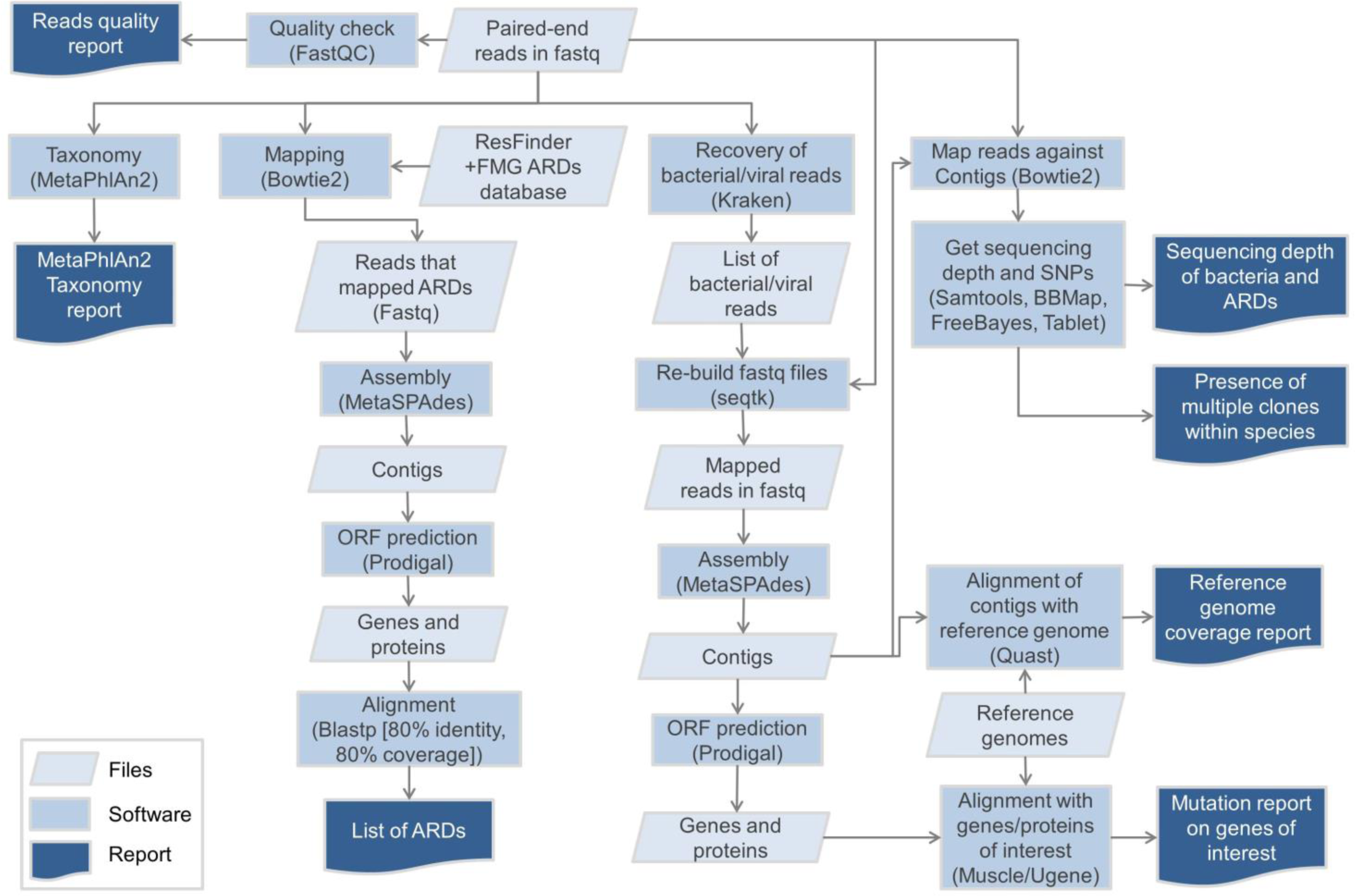
Bioinformatic analysis performed in this study. ARDs: antibiotic resistance determinants. Fastq: format for the files that embeds the read sequences 2 and their per-base quality score. FMG: functional metagenomics; ARDs: antibiotic resistance determinants; ORF: open reading frame.

## Results

### DNA extraction

We first extracted the samples for which the quantity of material exceeded or was equal to 1 mL (n=77), and the two swabs. We recovered more than 1 pg bacterial DNA mostly for samples that had grown at >100 CFUs (Supplementary Figure 1, panel A), while the concentration of human DNA did not seem to correlate with bacterial load (Supplementary Figure 1, panel B). Accordingly, the remaining samples, that had grown at least 100 CFUs (n=25), were submitted to extraction. In total, the DNA of 104 samples was extracted, from which 24 met the requirement to be sequenced (*i.e.* contained at least 1 pg/µL bacterial DNA, and less than 99% human DNA, Supplementary Table 1). Among the 24 samples and all throughout the manuscript, we will refer as monomicrobial (n=8) and polymicrobial (n=16) samples those which respectively yielded one and more than one bacterial species in culture.

**Table 1:** Characteristics of the 14 patients for whom 24 samples were sequenced. ASA: American Society of Anesthesiologists.

| Patient | Samples | Age | Gender | ASA score | Body mass index | Post-operative infection (type of surgery) | Delay between surgery and infection | Body site | Material involved |
| --- | --- | --- | --- | --- | --- | --- | --- | --- | --- |
| A | 2, 66, 28 | 51 | M | 2 | 30.4 | Yes (material) | <1 month | Ankle | Osteosynthesis |
| B | 4, 140 | 50 | F | 2 | 39.8 | No | NA | Clavicle | None |
| C | 19, 103, 104 | 54 | M | 2 | 24.1 | Yes (material) | <1 month | Toe | Osteosynthesis |
| D | 110 | 66 | M | 2 | 29.4 | Yes (material) | Between 1 and 3 months | Tibia | Osteosynthesis |
| E | 42 | 61 | F | 3 | 50.6 | Yes (material) | <1 month | Knee | Total knee prothesis |
| F | 46 | 63 | M | 2 | 18.0 | Yes (material) | <1 month | Mandible | Osteosynthesis |
| G | 59, 117, 136 | 69 | M | 2 | 25.5 | Yes (bone resection) | NA | Tibia | None |
| H | 90, 158 | 64 | F | 2 | 21.2 | No | NA | Sacrum | None |
| I | 108, 181 | 86 | F | 2 | 26.7 | Yes (material) | Between 1 and 3 months | Knee | Total knee prothesis |
| J | 121, 172 | 50 | F | 1 | 24.2 | No | NA | Tibia | None |
| K | 128 | 86 | F | 2 | 30.1 | No | >3 months | Knee | Osteosynthesis |
| L | 171 | 51 | M | 1 | 25.6 | Yes (material) | >3 months | Tibia | Osteosynthesis |
| M | 178 | 87 | F | 3 | 26.1 | Yes (material) | <1 month | Knee | Total knee prothesis |
| N | 184 | 60 | M | 3 | 34.3 | No | NA | Greater trochanter and ischiur | None |

### Bioinformatics

After trimming, we obtained a mean number of 10,046,084 reads per sample (range 4,128,425-14,549,687, Supplementary Table). With the Kraken classifier, the mean rate of classified reads (as bacteria, archea or virus) was 27.9% (range 1.8-85.7, Supplementary Table 1). Of note, the classification rate was correlated to the proportion of bacterial DNA as found by qPCR (Pearson’s correlation test, p<0.001, Supplementary Figure 2). The assembly of the classified reads with metaSPAdes yielded a mean number of contigs of 10,444 (range 3,087-18,513, Supplementary Table 1), for a mean total number of base pairs of 8.3M (range 2.9M-16.5M, Supplementary Table 1). Of note, the total number of base pairs of contigs was higher in polymicrobial samples than in the monomicrobial ones (respectively 9.7M vs. 5.5M, t test p <0.05, Supplementary Figure 3). The mean size of the contigs was 805 bp (median 369 bp, maximum 445,300 bp, Supplementary Figure 4).

### Identification of the pathogens

In monomicrobial samples (n=8, Table 2), 8/8 (100%) of the pathogens identified by culture were found by MetaPhlAn2, mostly at very high abundances (over 94.6%) at the exception of sample 46 in which *Streptococcus anginosus* was only found at a 2.2% abundance (Figure 2). In polymicrobial samples (n=16, Table 2), 55 bacterial species were found in culture 32 of which (58.2%) were found by MetaPhlAn2. At the genus level the match rate increased to 41/55 (74.5%). The presence of all bacteria found by culture in a given sample was confirmed by MetaPhlAn2 for 11/24 (45.8%) samples at the species level, including 3/16 (18.8%) for polymicrobial infections. At the genus level, 15/24 (62.5%) samples were in agreement with cultures, including 7/16 (43.8%) samples with polymicrobial infections.

**Figure 2:**
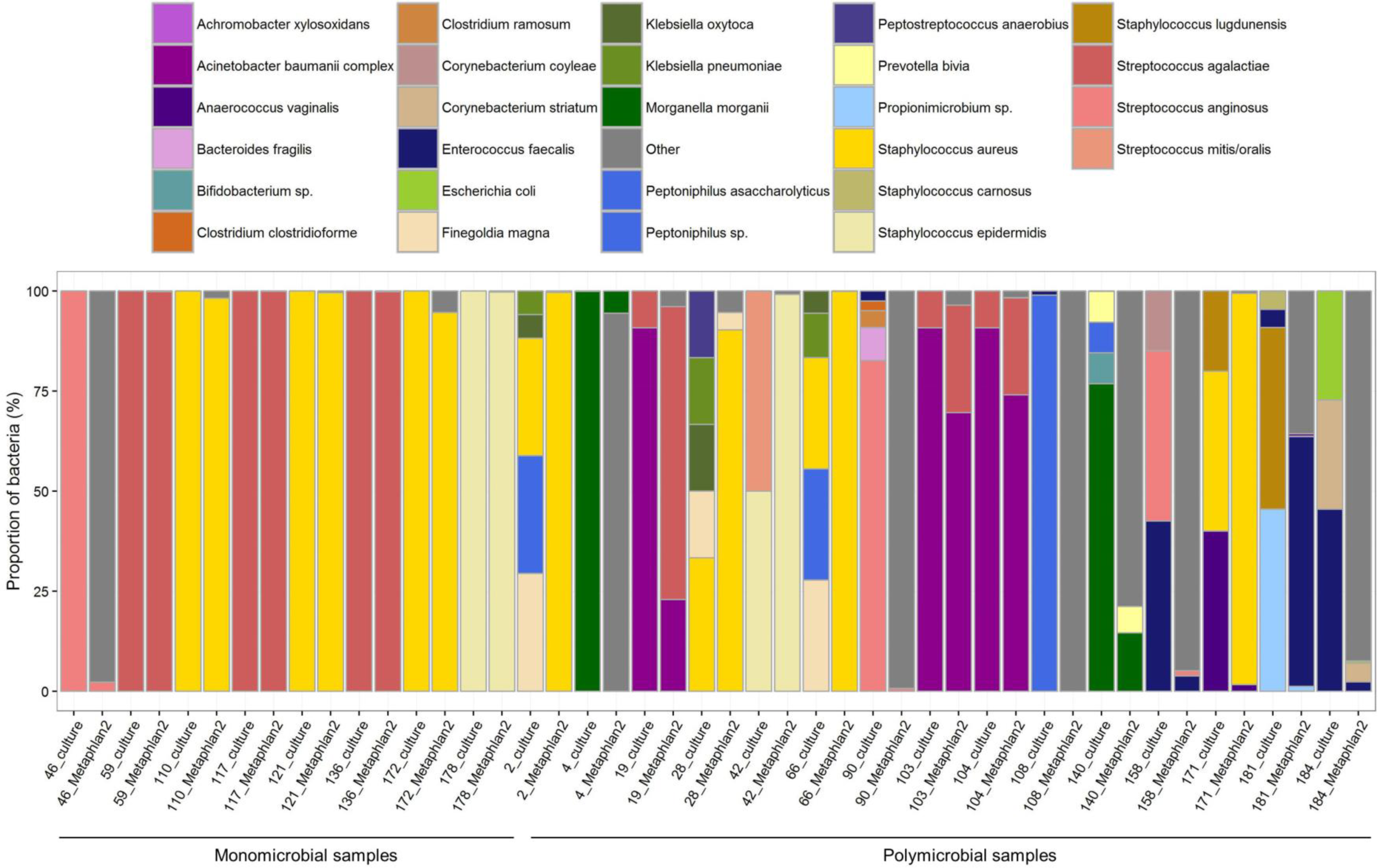
Proportions of the species recovered in culture and from reads (using MetaPhlAn2 [19]).

**Table 2:**
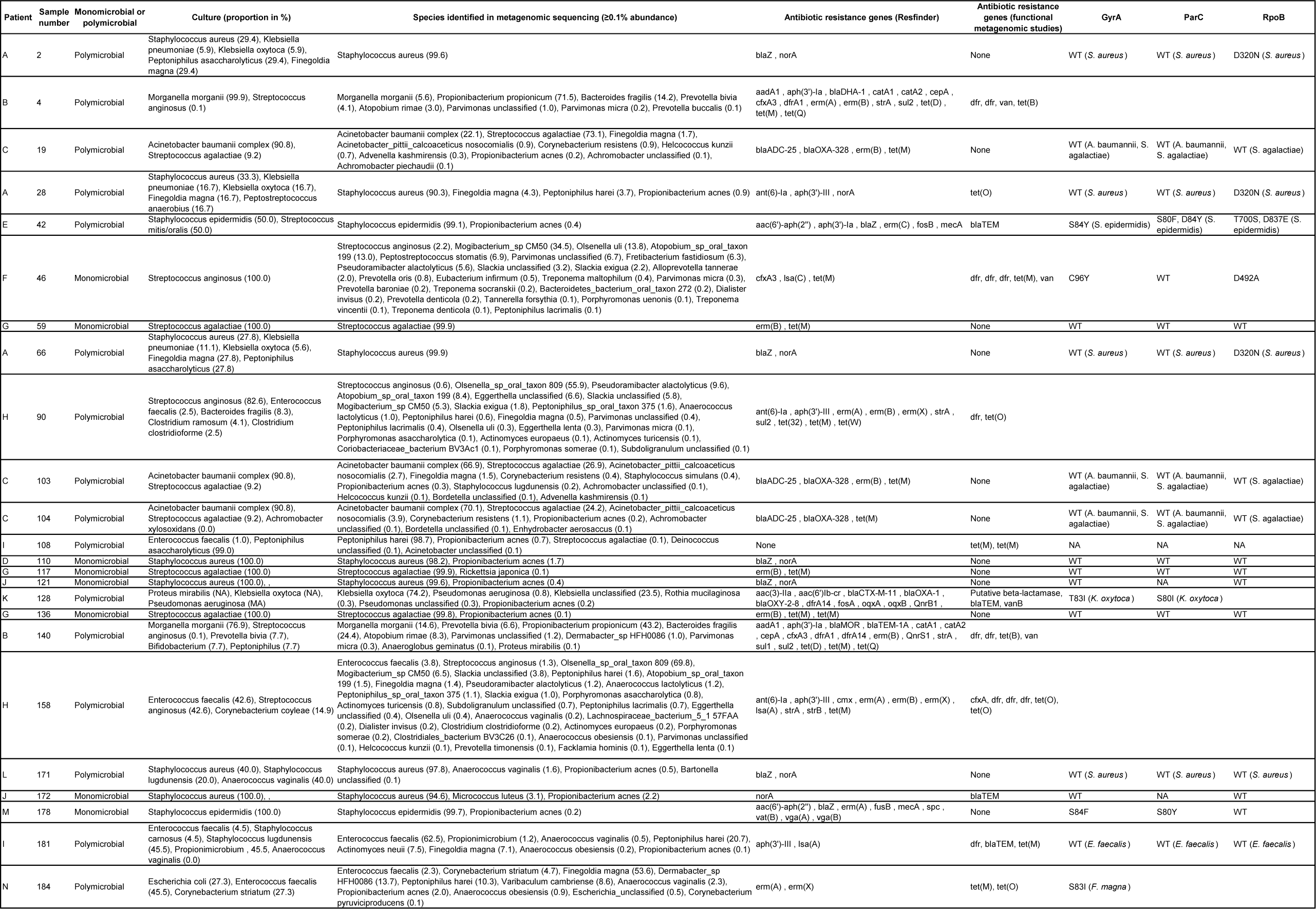
Description of the 24 samples sequenced in this study. WT: wild-type. NA: not assembled.

### Identification of other bacteria and possible contaminants

Apart from the bacteria that were found in culture (n=63) in the 24 positive samples, a total of 273 bacteria, not found in culture, were identified by MetaPhlAn2 (Figures 2 and 3). One (*Propionibacterium acnes*) was found in 20/24 samples (supplementary Figures 5 and 6). Moreover, the abundance of *P. acnes* in samples was negatively correlated to their total DNA concentration sample (Supplementary Figure 5), supporting that *P. acnes* was a contaminant in this study [25,26]. For other species, such correlation could not be tested because of their low occurrence in samples. We identified 66 likely contaminants (Figure 3, Supplementary Table 2), some being commonly found in culture (such as *Micrococcus luteus*) or as reagents contaminants [27]. Others were unexpected such as *Borrelia sp.* (samples 103, 104, 108 and 110) or *Rickettsia japonica* (sample 117). Still, the taxonomic assignment of the contigs did not confirm the presence of those species and manual blastn of reads against the NCBI nr database supported that they were likely *in silico* contaminants (data not shown) [27,28]. Besides, we identified 25 species that could be due to a misclassification of reads to closely related bacteria, such as in samples 184 (where *Corynebacterium striatum* was found in culture, and some metagenomic reads were identified as *Dermabacter* sp. and *Corynebacterium pyruviciproducens*), samples from patient C (19, 103, 104 where *Acinetobacter baumannii* and *Achromobacter xylosoxidans* were found in culture, and some reads were identified as from other *Acinetobacter* spp., *Achromobacter* spp., or *Advenella kashmirensis*, a bacterium close to *Achromobacter*) (Supplementary Table 2). Hence, a total of 182 bacteria not recovered in culture and not acknowledged as contaminants were identified in metagenomic sequencing. For one sample that was monomicrobial in culture (sample 46, that yielded *S. anginosus*), 38 other species were identified by metagenomics. Interestingly, these species appeared to be commonly found in the oropharyngeal microbiota, which was consistent with the site of the infection (mandible). In polymicrobial samples such as samples 4 and 140 (patient B), 90 and 158 (patient H), 108 and 181 (patient I), metagenomic sequencing identified several more anaerobic bacteria (range 3-40, see Supplementary Table 2, in consistence with the sporadic isolation of such bacteria in the routine culture of these of samples. In both samples 4 and 140 from patient B, the most abundant species was *Propionibacterium propionicum* (respective abundances of 71.5% and 43.2%) that was not found in culture. Arguments in contradiction with *P. propionicum* being a contaminant in these samples are that the species found in other samples was *P. acnes*, and that the abundance of *P. propionicum* was high (supplementary Figure 6) whereas the abundance of *P. acnes* was low in the samples where it was identified.

**Figure 3:**
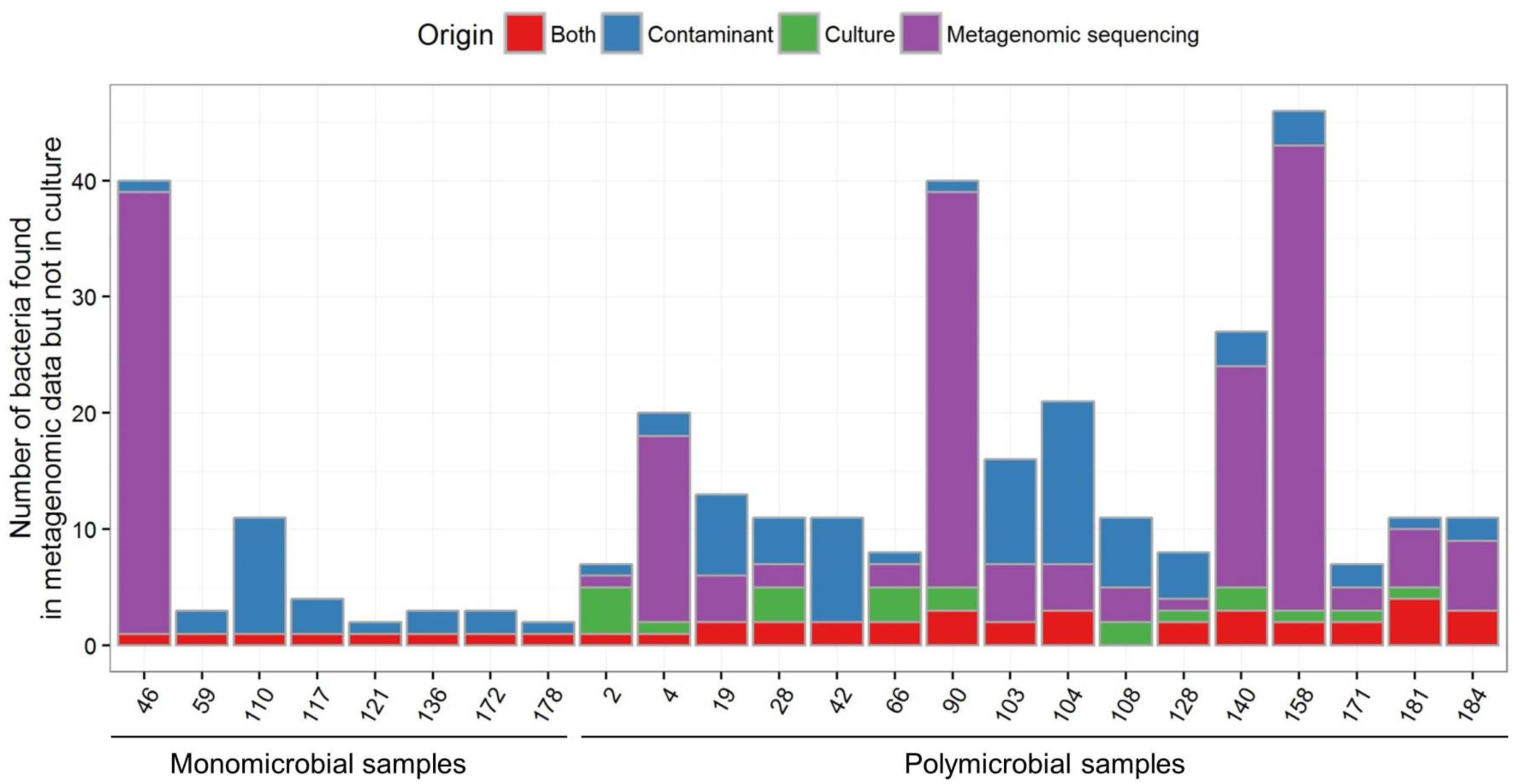
Distribution of the number of species found both in culture and metagenomic sequencing, only in culture and only metagenomic sequencing (in this case, putative pathogenic species and contaminants/misclassifications are depicted apart). All species identified by MetaPhlAn2 were considered (not only those above 0.1% abundance).

### Identification of clones within species

Based on this assumption that in case of multiple clones within one species, the SNVs would be homogeneously distributed along the contigs (see Supplementary methods and Supplementary Figure 7), we found polyclonal populations for 29 of the 74 (39.2%) bacterial species that were tested. Among the bacteria that were found in culture and that were tested (n=32), 8 (25%) displayed a polyclonal population: *Morganella morganii* (samples 4 and 140), *Streptococcus agalactiae* (samples 103 and 117), *Staphylococcus aureus* (samples 28 and 110), *S. anginosus* (sample 158) and *Pseudomonas aeruginosa* (sample 128). Moreover, we observed that for *M. morganii* (samples 4 and 140), no mutations on the topoisomerases of M. morganii were found in the sample 140, while in sample 4 the Ser83Ile and Ser84Ile were found the in GyrA and ParC, respectively. This suggests that one population of *M. morganii* was susceptible to fluoroquinolones and the other was not. In culture though, only the fluoroquinolone resistant clone was found.

### Antibiotic resistance determinants, linkage with the host and inference of antibiotic susceptibility

A total of 151 ARDs (61 unique) were identified from the 24 samples (range 2-22, Table 2). The most frequent ARD families were beta-lactamases (n=30), Tet(M) (n=26), Erm (n=18) and Dfr (n=16). For monomicrobial samples, we assumed that the ARDs identified by metagenomics were expressed by the bacterium that was recovered in culture. Considering together (*i*) the antibiotic class the ARDs usually confer resistance to, (*ii*) the antibiotic susceptibility of wild-type species and (*iii*) the analysis of the sequence of specific genes (*gyrA*, *parC*, *rpoB*), we could infer a *in silico* susceptibility in agreement with the phenotypic susceptibility in 94.1% (111/118) cases (Figure 4 and supplementary tables, a case being defined as the susceptibility testing of one antibiotic for one sample). Of note, the six major errors (overprediction of resistance as compared to culture) originated from sample 46 where anaerobic bacteria and likely associated ARDs were found in metagenomic sequencing but not in culture. For polymicrobial samples, as we could not rely on the depth of sequencing of ARDs and bacterial contigs to infer some connections (Supplementary Figure 8), we separately considered the ARDs and the bacteria found in the sample (Supplementary Table 2). Accordingly, we inferred a correct susceptibility in 76.5% (192/251) cases. Very major errors mostly occurred because some bacteria with specific resistance patterns were not detected in sequencing (Supplementary Table 2). Conversely and along with the observations with monomicrobial samples, most major errors occurred because some bacteria and ARDs were found in sequencing but not in culture. Of note, the prediction of susceptibility to fluoroquinolones was correct in 100% (24/24) samples

**Figure 4:**
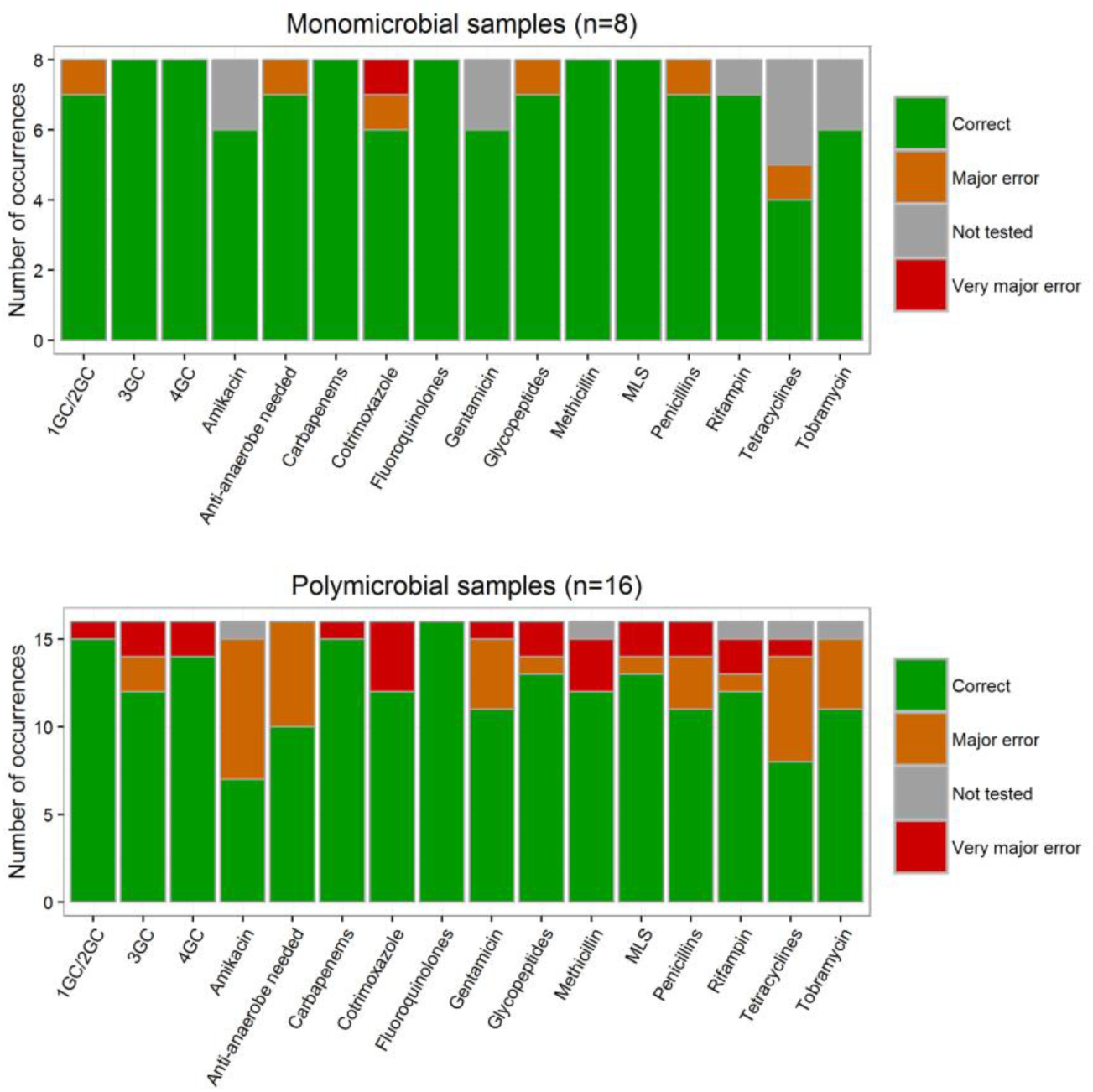
Antibiotic susceptibility inference from metagenomic data compared to culture and conventional antibiotic susceptibility testing (gold standard). 1GC/2GC, 3GC, 4GC: 1^st^, 2^nd^, 3^rd^ and 4^th^ generation cephalosporins, respectively. MLS: macrolides, lincosamides, streptogramines. Correct: metagenomic result consistent with the result given by conventional methods. Very major error: metagenomic data did not predict antibiotic resistance while at least one bacteria identified by conventional methods was resistant to this antibiotic. Major error: metagenomic data predicted antibiotic resistance while all the bacteria identified by conventional methods were susceptible. Not tested: no molecule from the antibiotic class was tested with conventional methods.

### Influence of downsizing the samples to 1M reads

We ran the same pipeline analysis onto the 24 samples downsized at 1M reads. We observed that the taxonomic distribution did not apparently change for the most abundant species (Supplementary Figure 9), but the mean genome coverage of the main pathogen was lower in the downsized group than in the full-reads group (3.9% vs. 8.9%, Student paired test p<0.001, Supplementary Figure 10). Also, only 86 ARDs were found after downsizing while 151 were detected before (Student paired test p<0.001, Supplementary Figure 9). Of note, the impact of downsizing was observed in both monomicrobial and polymicrobial samples (Supplementary Figures 11 and 12).

## Discussion

The main result of this study is that we showed that metagenomic sequencing could be a potential tool in the diagnostic of BJI. Indeed for monomicrobial infections, the pathogen was identified in 100% (8/8) samples and the antibiotic susceptibility prediction was successful in 94.1% (111/128) cases. In case of polymicrobial samples, the high abundance of several bacteria (mostly anaerobes) did occasionally prevent from the correct identification of the pathogens and their antibiotic susceptibility profiles. Accordingly, our findings support that currently, metagenomic sequencing of BJI samples could not replace conventional methods based on culture due to the limitations encountered when several bacterial populations are present in the samples, but rather be performed in support.

Interestingly, metagenomic sequencing yielded in some ways more information than culture. First, metagenomic sequencing identified many more bacterial species than culture. Besides likely contaminants, some bacteria were probably true positive that were not detected by culture and may not have been targeted by the selected antibiotic regimen. Second, we could observe within species at least two clonal populations, which could differ in their susceptibility to antibiotics as we observed for fluoroquinolones in *M. morganii*. In all, using metagenomic data could help to tailor the antibiotic regimen for the treatment of BJI, and the added-value of clinical metagenomics in BJI should now be assessed.

However, there are several hindrances to the application of metagenomic sequencing to BJI samples. First, we could only sequence 24 out of 179 samples, due to a low amount of bacterial DNA that could be recovered from the samples. This is the main limitation of this study as it reduced the diversity of clinical situations that we could address. Nonetheless, the samples from this study have been frozen and thawed, which decays bacteria and releases DNA. As the DNA extraction method we used eliminates free DNA after lysing eukaryotic cells, it is likely that we could have sequenced more samples if they would not have been frozen. This said, recovering enough bacterial DNA (in terms of quantity and proportion with respect to human DNA) remains challenging. Also, the high cost of NGS currently prevents its routine application. We tested the impact of a lower depth of sequencing and showed that despite the taxonomic profiles of the bacterial populations were similar, the inference of antibiotic susceptibility was less accurate due to a lower recovery of genes involved in antibiotic resistance. Our results suggest that clinical metagenomics should indeed benefit from the highest depth of sequencing. Another limitation of the study is that we did not concomitantly sequence a negative control to identify the putative contaminants that would originate from the sample process. Some contaminants have been identified in studies using 16S rDNA amplifications [27], but some of them are also met as BJI pathogens (e.g. *P. acnes*). A solution to this issue would be to include a negative control for every run or at least when new batches of kits are used, and to subtract from the clinical samples the bacteria found in the negative control based on their abundance [29].

Besides, our observations suggest that clinical metagenomics will soon require, as for clinical microbiology, a specific expertise combining clinical, biological and bioinformatic skills in order to infer clinically relevant results from metagenomic data. In this perspective, the development of clinical metagenomics will need the definition of quality standards, e.g. what is the sufficient genome coverage for a given bacterium to consider that its antibiotic susceptibility profile can be likely inferred. In the long term, algorithms should be built to provide clinicians with clear data and robust algorithms to support clinical decisions.

In conclusion, we showed that metagenomic sequencing of BJI samples was a potential tool to support conventional methods. In this perspective, its main limitations (DNA extraction, cost and data management) should be tackled, and the clinical benefit provided by clinical metagenomics should now be assessed in a prospective fashion.

## Acknowledgements

None.

## Supplementary Figures

**Supplementary Figure 1:**
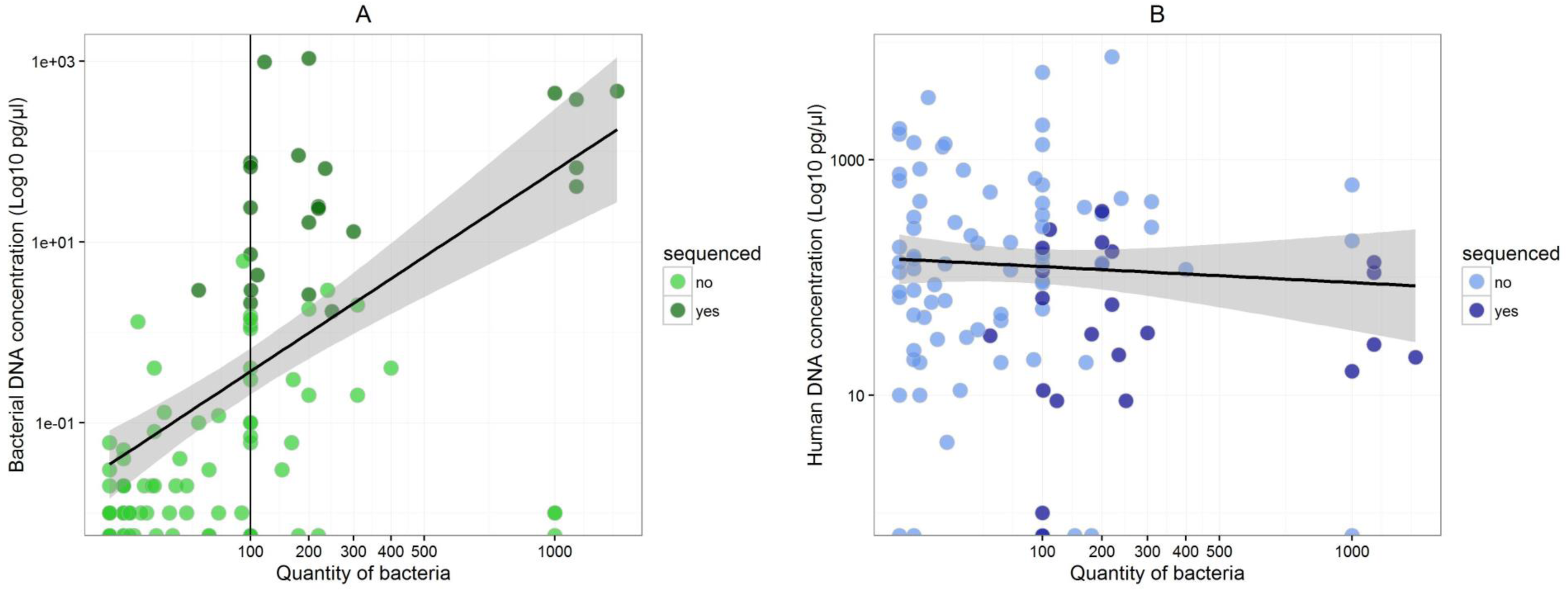
Total quantity of bacteria obtained in culture and concentrations of bacterial (A) and human DNA (B) for 102 samples for which DNA was extracted. DNA concentrations in DNA extracts were determined by qPCR (see methods). The shaded grey area depicts the 95% confidence interval around the linear regression line. The X-axis is square-root transformed for visibility purposes. One sequenced sample is missing (sample 128) because the bacterial concentrations were missing. Eventually, 19 and 7 samples with null bacterial and human DNA concentrations are not shown in the panel A and B, respectively.

**Supplementary Figure 2:**
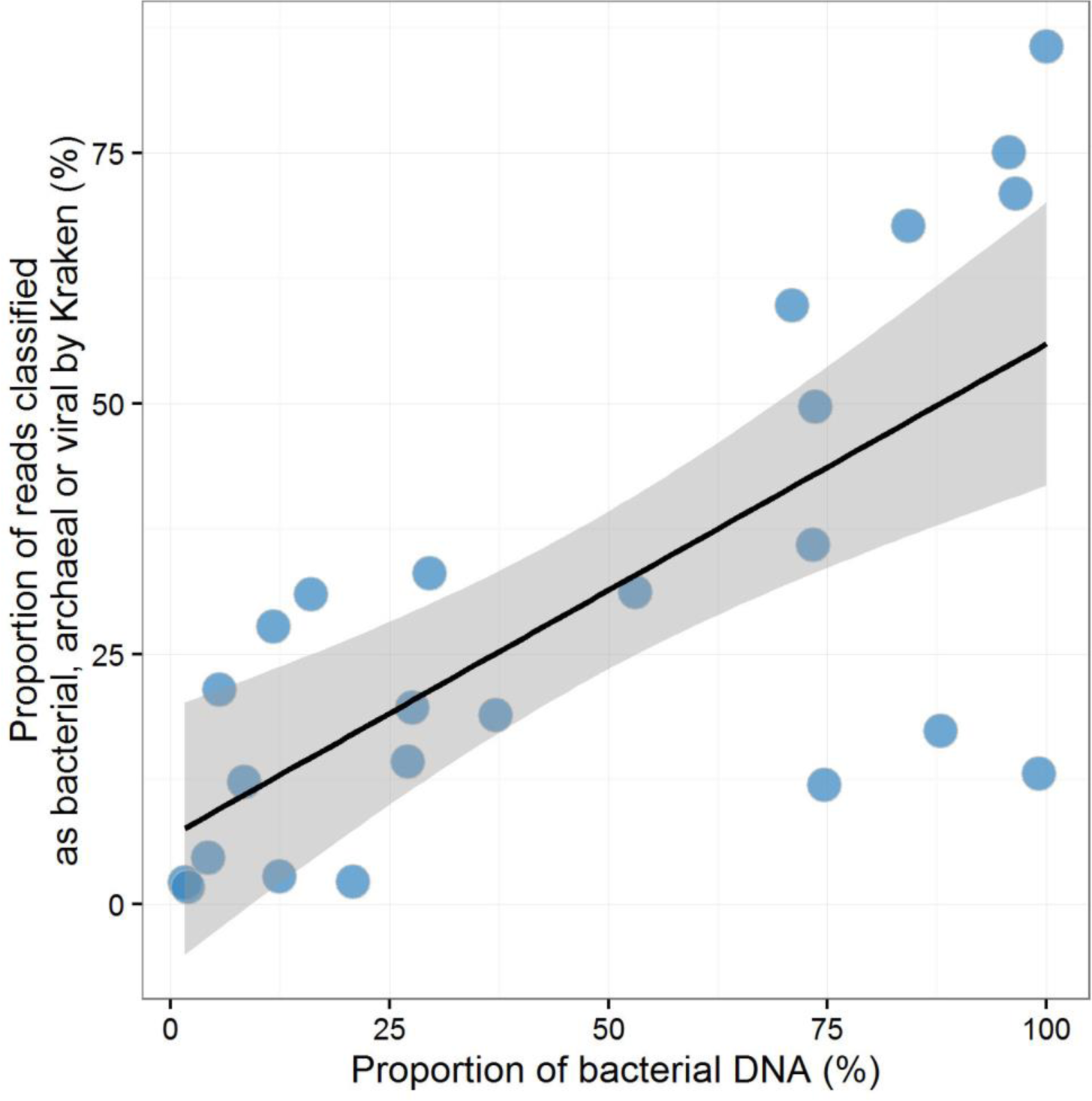
Proportion of bacterial DNA as determined by qPCR (the proportion being calculated as the percentage of bacterial DNA on the total DNA [bacterial and human]) on the X-axis, and proportion of the quality-filtered reads classified as bacterial, archaeal or viral by the Kraken classifier. The shaded grey area depicts the 95% confidence interval around the linear regression line.

**Supplementary Figure 3:**
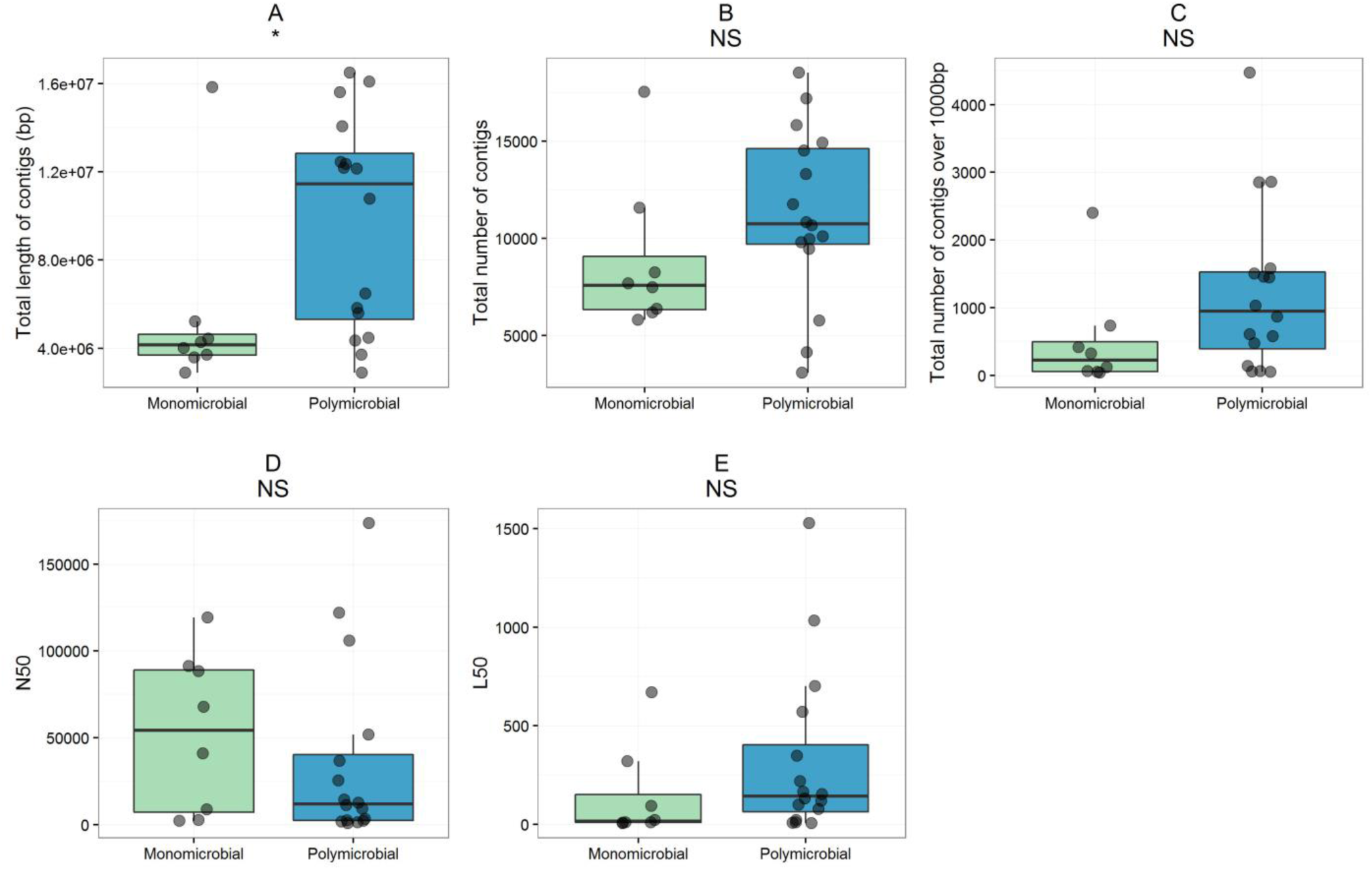
Boxplots of the the length of summed contigs (A), the total number of contigs (B), the total number of contigs exceeding 1000bp (C), the N50 (D), the L50 (E) according to the number of bacteria recovered in culture (monomicrobial or polymicrobial). The contigs were obtained by the assembly by MetaSPAdes [20] of reads classified by Kraken [30]. The parameters showed on this figure were obtained by Quast [31]. *: p<0.05; NS: not significant. The boxplot limits represents (from bottom to roof) the 25^th^, 50^th^ (median) and 75^th^ percentiles.

**Supplementary Figure 4:**
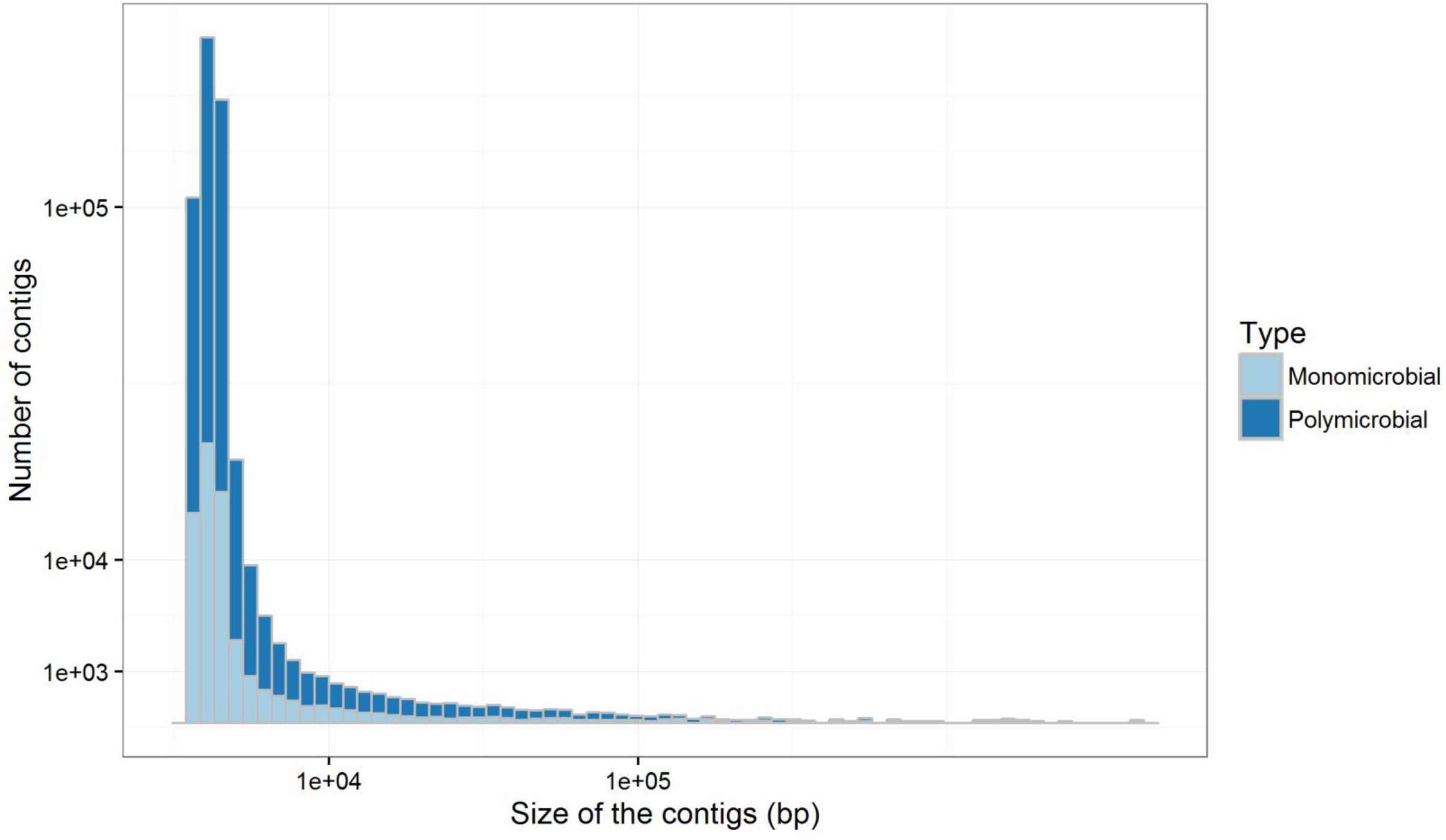
Distribution of the size of the contigs for all the samples (n=24). Both Y-axis and X-axis are square-root transformed.

**Supplementary Figure 5:**
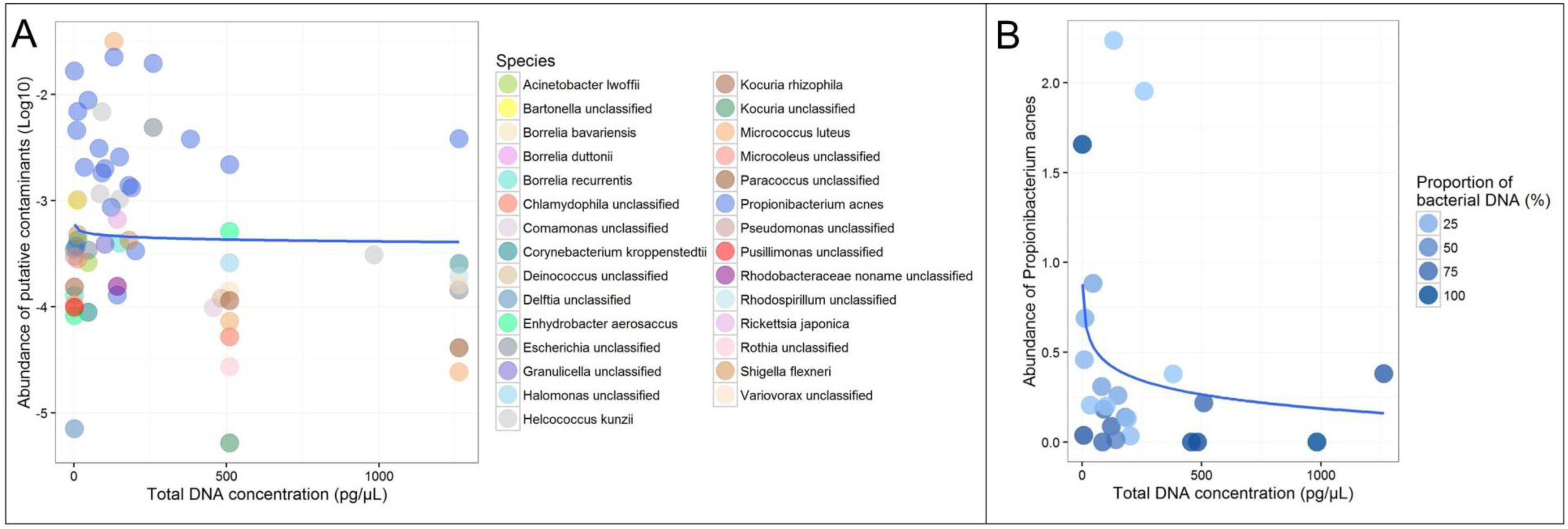
(A) Dot plot of the abundance of the species (Log10) that were considered as contaminants in this study (see supplementary Table 1) along with the total DNA (sum of bacterial and human DNA) concentrations (pg/µL). The blue line depicts the linear regression between the abundances of the species (Log10) and the Log10 of the DNA concentrations. (B) Dot plot of the abundance of Propionibacterium acnes (%) along with the total DNA (sum 3 of bacterial and human DNA) concentrations (pg/µL). Samples with an abundance of 0 are also showed. The blue line depicts the linear regression between 4 the abundances of the species and the Log10 of the DNA concentrations.

**Supplementary Figure 6:**
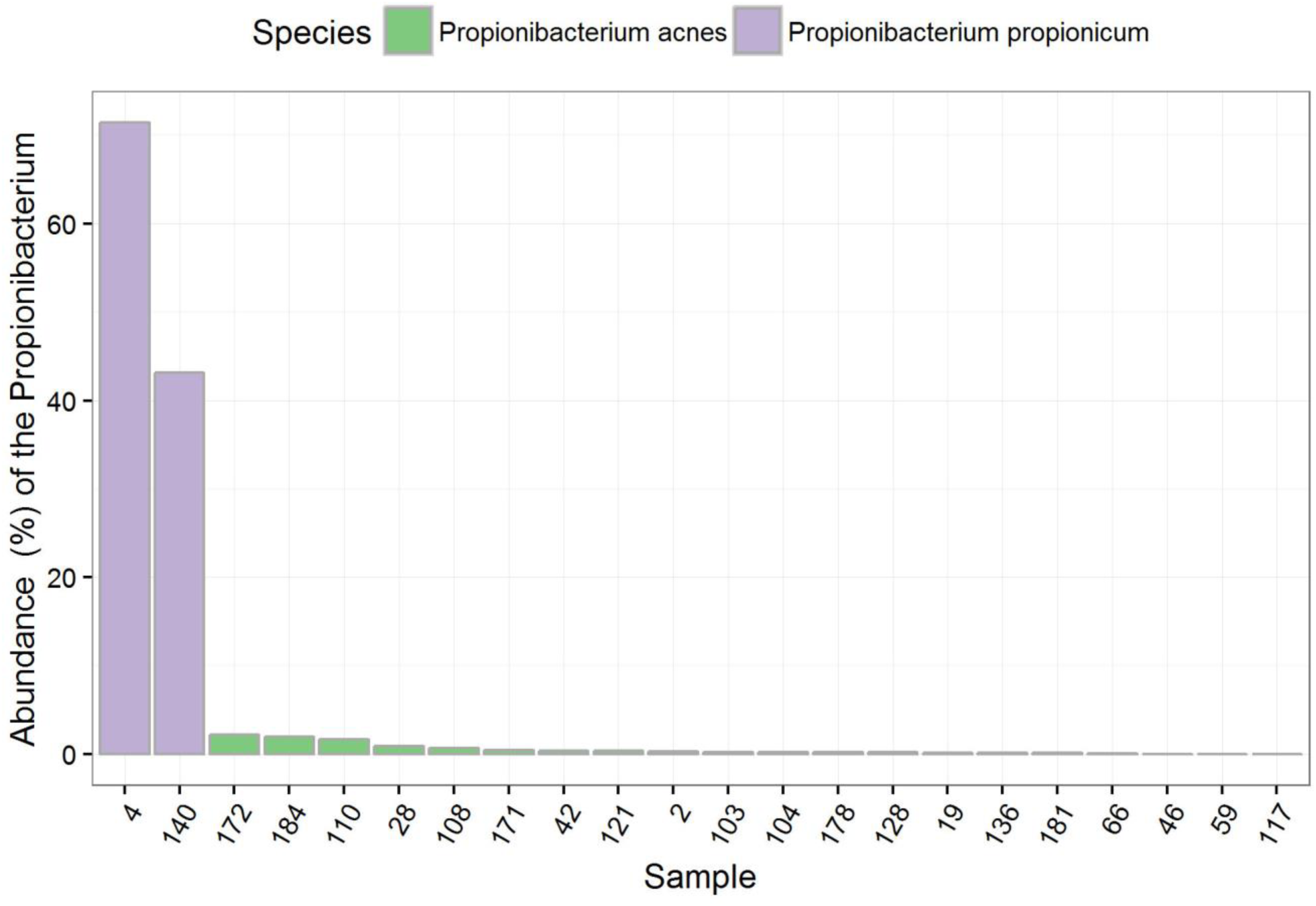
Distribution of the abundances (%) of *Propionibacterium propionicum* (purple) and *Propionibacterium acnes* (green) in the 22 samples where they were identified.

**Supplementary Figure 7:**
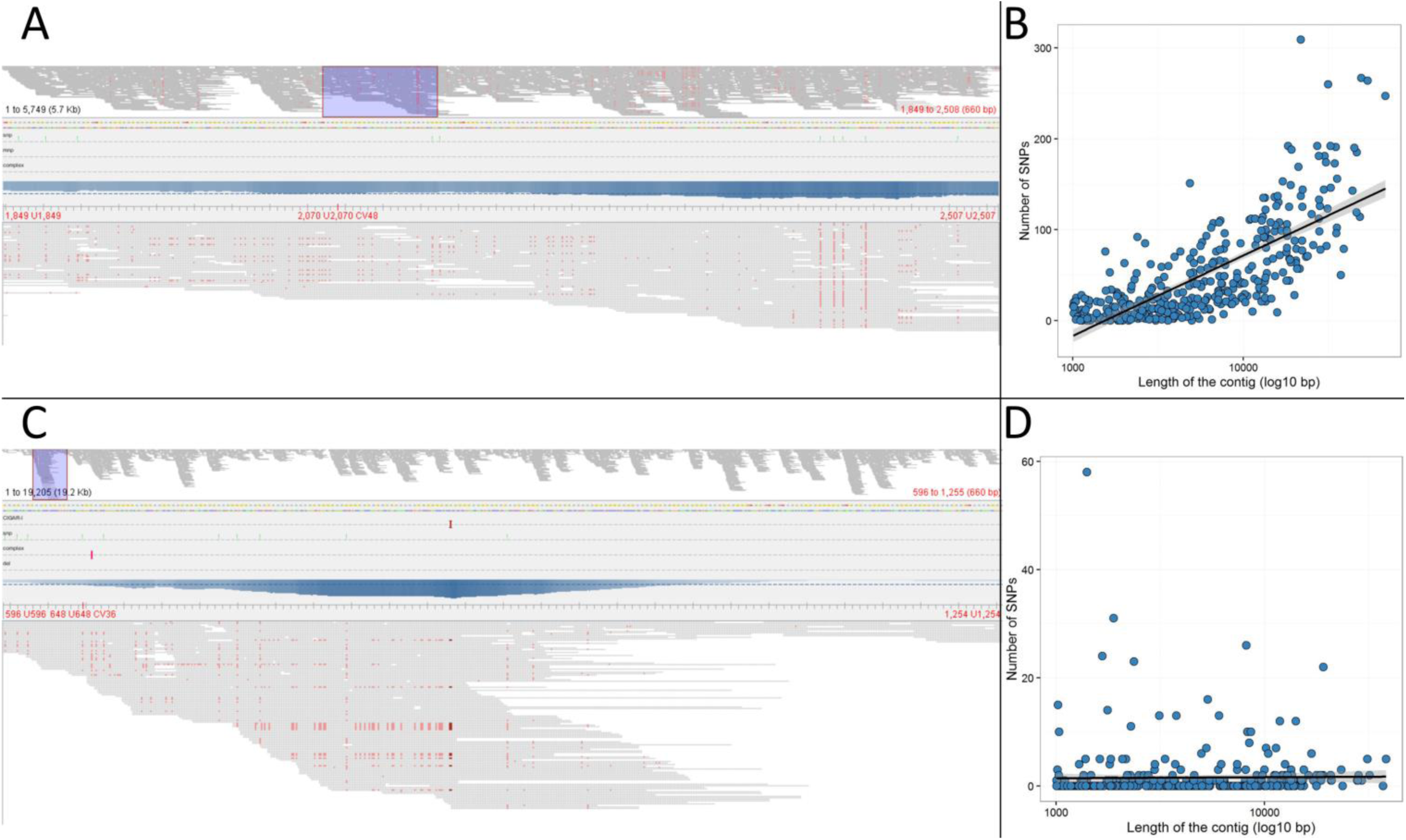
Examples of the assessment of a polyclonal population within species. A: The Tablet view of a 5,749 bp contig from *Morganella morganii* in sample 4. B: The dot-plot of the number of SNVs and the size of the contigs from *Morganella morganii* in sample 4 (Pearson’s correlation test p<0.001). C: The Tablet view of a 19,205 bp contig from Staphylococcus aureus in sample 121. In this specie, the SNVs are concentrated in some regions (such as the one showed) and not homogeneously scattered. D: The dot-plot of the number of SNVs and the size of the contigs from *Staphylococcus* aureus in sample 121 (Pearson’s correlation test p=0.4).

**Supplementary Figure 8:**
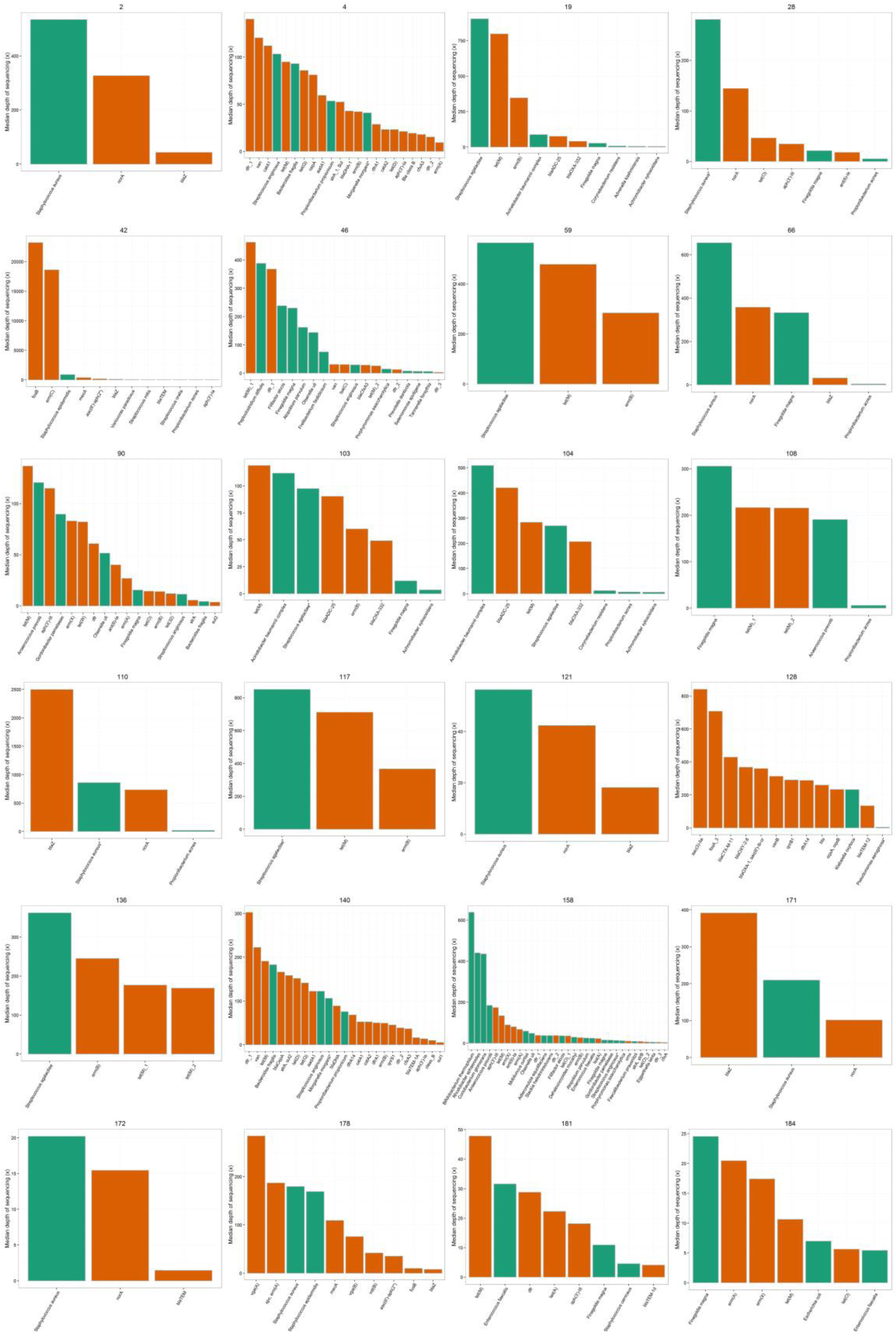
Bar plot of the median depth of sequencing (expressed in ×) of the contigs of the bacterial species (in green) and depth of sequencing of the ARDs (in orange) found in the sample.

**Supplementary Figure 9:**
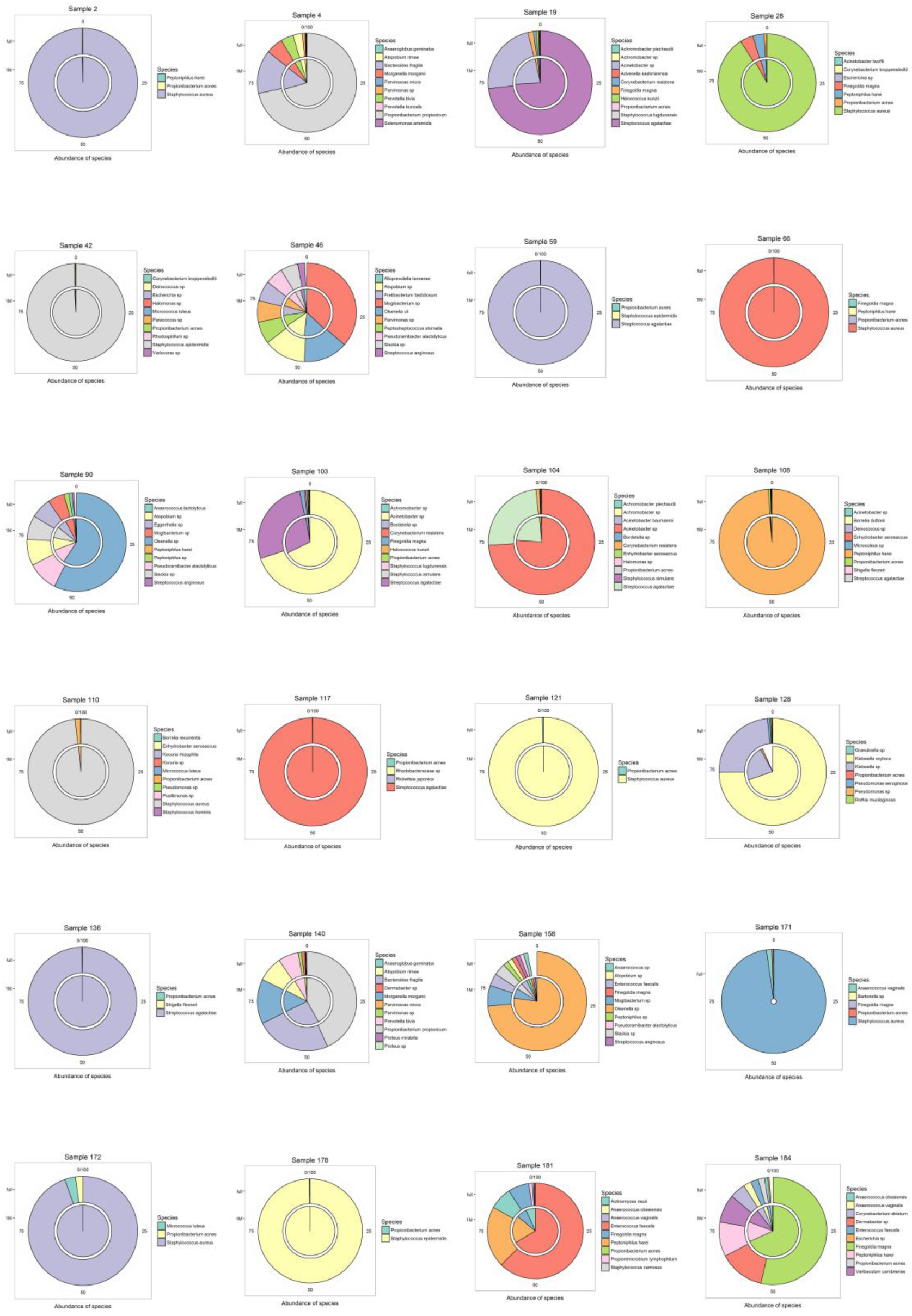
Influence of the downsizing to 1M reads on the taxonomic classification of reads by MetaPhlAn2. The outer and the inner circles represent the distribution of the main species with no and 1M downsize, respectively.

**Supplementary Figure 10:**
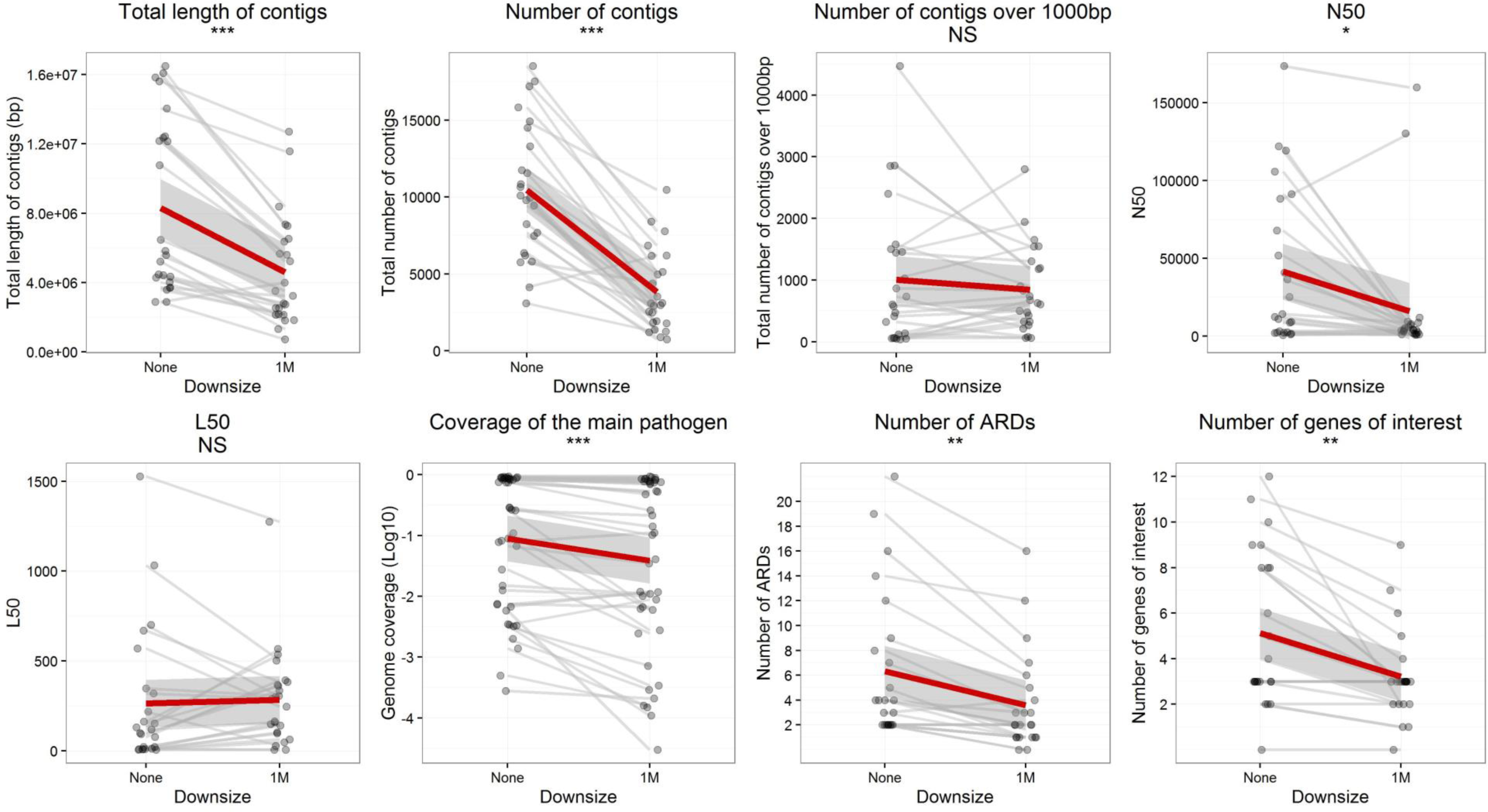
Effect of downsizing to 1M reads on the assembly performances for all samples (n=24). Paired t-tests were performed. ARDs: antibiotic resistance determinants. The genes of interest included *gyrA*, *parC* and *rpoB* (with sizes over 80% of the reference genes). ***: p<0.001; **:p<0.01; 2 NS: not significant. The shaded grey area depicts the 95% confidence interval around the linear regression line.

**Supplementary Figure 11:**
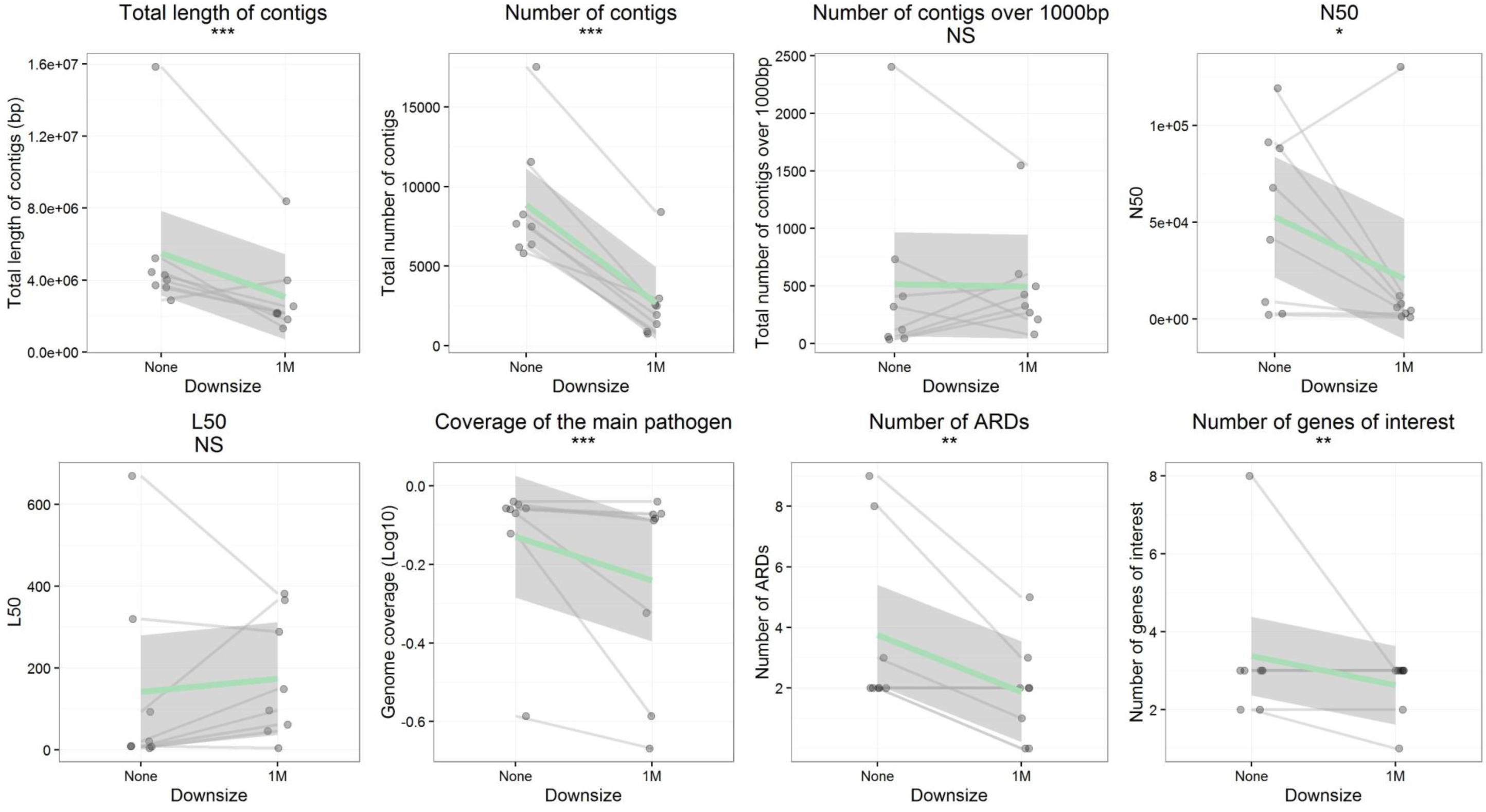
Effect of downsizing to 1M reads on the assembly performances for monomicrobial samples (n=8). Paired t-tests were performed. ARDs: antibiotic resistance determinants. The genes of interest included *gyrA*, *parC* and *rpoB* (with sizes over 80% of the reference genes). ***: p<0.001; **:p<0.01; NS: not significant. The shaded grey area depicts the 95% confidence interval around the linear regression line.

**Supplementary Figure 12:**
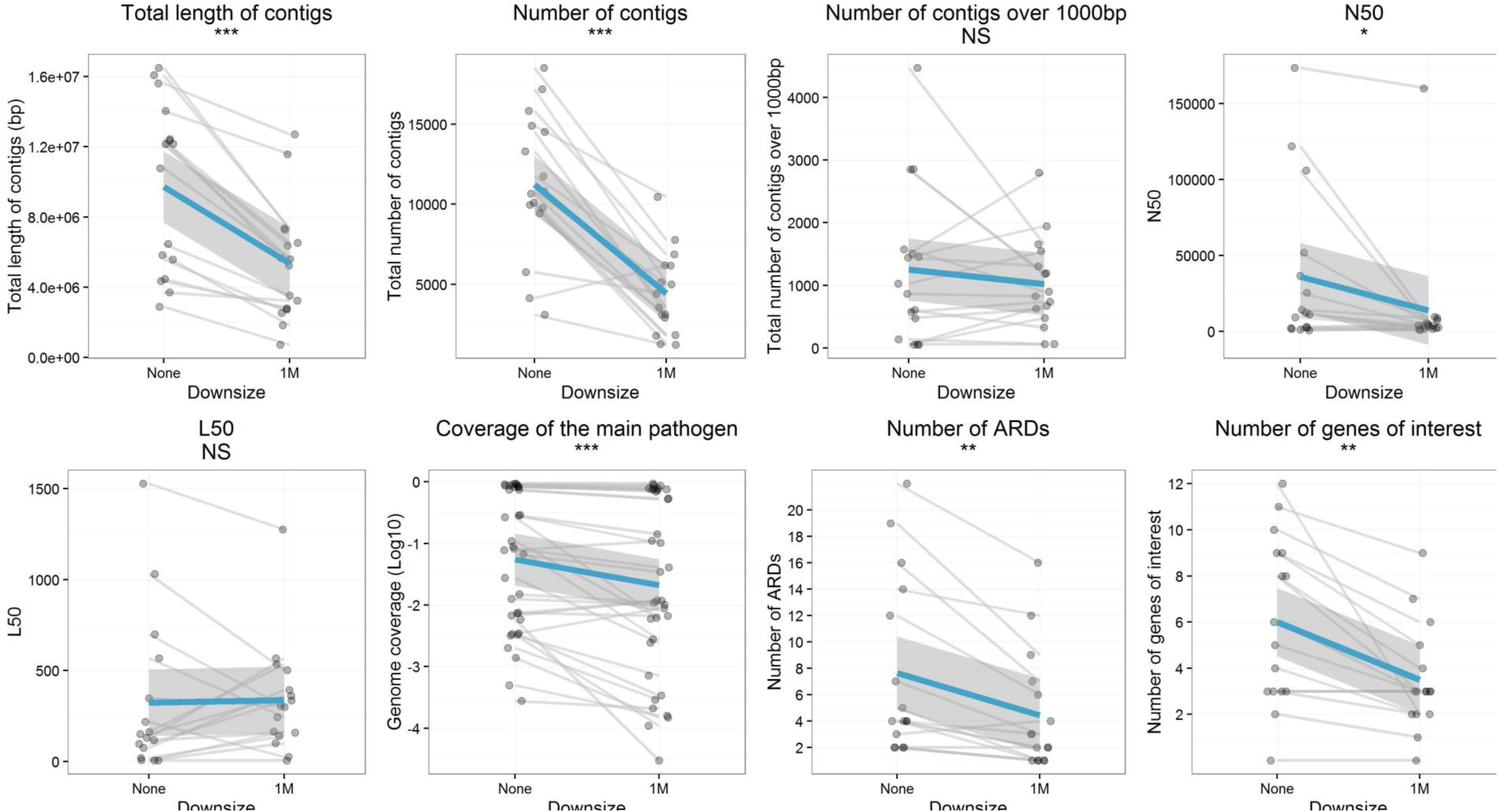
Effect of downsizing to 1M reads on the assembly performances for polymicrobial samples (n=16). Paired t-tests were performed. ARDs: antibiotic resistance determinants. The genes of interest included *gyrA*, *parC* and *rpoB* (with sizes over 80% of the reference genes). ***: p<0.001; **:p<0.01; NS: not significant. The shaded grey area depicts the 95% confidence interval around the linear regression line.

## Supplementary Tables legend

**Supplementary Table 1**: Summary of the sequencing and assembly results for the 24 samples. N50 is the median contig size of the metagenomic assembly. L50 is the number of contigs that accounts for more than 50% of the metagenomic assembly.

**Supplementary Table 2**: Inference of antibiotic susceptibility from metagenomic data and antibiotic susceptibility of the bacteria found in culture.

